# Age-dependent modulation of the excitability of layer V pyramidal neurons by dopamine D1 receptors in mice’s primary motor cortex

**DOI:** 10.1101/2023.06.06.543824

**Authors:** Valentin Plateau, Jérôme Baufreton, Morgane Le Bon-Jégo

## Abstract

The primary motor cortex (M1) receives dopaminergic (DAergic) projections from the midbrain which play a key role in modulating motor and cognitive processes, such as motor skill learning. However, little is known at the level of individual neurons about how dopamine (DA) and its receptors modulate the intrinsic properties of the different neuronal subpopulations in M1 and if this modulation depends on age. Using immunohistochemistry, we first mapped the cells expressing the DA D1 receptor across the different layers in M1, and quantified the number of pyramidal neurons (PNs) expressing the D1 receptor in the different layers, in young and adult mice. This work reveals that the spatial distribution and the molecular profile of D1 receptor-expressing neurons across M1 layers do not change with age. Then, combining whole-cell patch-clamp recordings and pharmacology, we explored *ex vivo* in young and adult mice the impact of activation or blockade of D1 receptors on PN intrinsic properties and identified a distinct modulation of intrinsic electrical properties of layer V PNs by D1 receptors depending on the age of the animal.

**Highlights:**

- The laminar distribution in M1 of cells expressing the dopamine D1 receptor is similar in young and adult mice
- Most of D1R-expressing cells in M1 also express Satb2
- D1R activation increases M1 layer V pyramidal neurons’ excitability both in young and adult mice
- The effect of D1R blockade on M1 layer V pyramidal neurons’ excitability differs between young and adult mice

## Introduction

Dopamine (DA) is a neuromodulator playing a key role in numerous physiological functions, such as cognitive (Nieoullon, 2002; Chudasama and Robbins, 2004; Floresco, 2013), reward (Yokel and Wise, 1975; Botvinick and Braver, 2015; Michely et al., 2020) and motor processes (Ungerstedt et al., 1969; Salamone, 1992; Alm, 2021). The dopaminergic (DAergic) system is of high importance and the consequences of its dysregulation are best illustrated by the diseases it triggers, notably Parkinson’s disease (Bernheimer and Hornykiewicz, 1965; Bernheimer et al., 1973; Albin et al., 1989; Nambu et al., 2015), schizophrenia (Davis et al., 1991; Brisch et al., 2014; Grace, 2016) and depression (Brown and Gershon, 1993; Grace, 2016). Thus, the role of DA in the striatum and the prefrontal cortex has been well documented (Robbins and Everitt, 1992; Diamond, 1996; D’Ardenne et al., 2012; Ott and Nieder, 2019; Valjent et al., 2019). However, even if to a lesser extent, the primary motor cortex (M1) also received DAergic innervation (Descarries et al., 1987; Gaspar et al., 1991), which comes from midbrain DAergic neurons (Hosp et al., 2011). M1 is involved in motor learning and it has been described that learning sophisticated motor sequences such as skill-reaching behavior relies upon DA-dependent structural and synaptic plasticity in M1 (Hosp et al., 2009, 2011; Guo et al., 2015). Although the architecture of the DA system within M1 has been well characterized anatomically in rodents (Descarries et al., 1987; Vitrac et al., 2014; Hosp et al., 2015), the level of understanding of DA action in M1 is rather macroscopic (Cousineau et al., 2022), monitoring global changes at the level of M1. Besides, its functional significance remains discussed, in part because the precise location of DA receptors and the modulation exerted by these receptors at the level of individual neurons are poorly understood and sometimes inconsistent.

DA activates two main classes of receptors, the D1-like and the D2-like family which are both present in M1 (Dawson et al., 1986; Lidow et al., 1989; Weiner et al., 1991; Gaspar et al., 1995). In M1, it has been shown that pyramidal neurons (PNs) and interneurons express D1 and/or D2 DA receptors (Gaspar et al., 1995; Vitrac et al., 2014; Cousineau et al., 2020; Swanson et al., 2021). Based on their axonal projection, PNs can be divided into 3 major classes in M1, the pyramidal tract neurons (PT) which express the transcription factor Ctip2 (also known as Bcl11b), the intra-telencephalic neurons (IT) which express Satb2 and the cortico-thalamic (CT) neurons mainly located in layer VI (Arlotta et al., 2005; Molnár and Cheung, 2006; Alcamo et al., 2008; Britanova et al., 2008; Shepherd, 2013; Digilio et al., 2015). Some studies investigated the effect of DA receptors on the intrinsic properties of M1 neurons (Parr-Brownlie, 2005; Vitrac et al., 2014; Cousineau et al., 2020), but the effects observed diverge (Parr-Brownlie, 2005; Vitrac et al., 2014; Cousineau et al., 2020; Swanson et al., 2021, for review see Cousineau et al., 2022), presumably because of the neuronal diversity, the different experimental conditions (*in vivo* versus *ex vivo*), the animal species (rats versus mice) and also the age of the animals. Indeed, age is a critical variable as developmental changes continue to occur far beyond the first postnatal weeks.

This study aimed to fill the gap in our understanding of how D1 receptors are expressed in the different layers of M1, and how they impact the intrinsic properties of PNs in the layer V of M1, in young and adult mice. Using the D1-GFP transgenic mouse line, we first mapped in young (P16-P25 old) and adult mice (6-12 weeks old) neurons expressing the D1 receptors according to M1 layers and pyramidal neuronal markers they express (Ctip2 and Satb2). Then using whole-cell patch-clamp recordings coupled to pharmacology, we investigated *ex vivo* in layer V how activation and blockade of the D1 receptor in M1 modulate PNs intrinsic properties in young and adult animals. This work reveals an age-dependent modulation of the excitability of M1 layer V PNs by D1 receptors.

### Experimental procedures Animals

All experiments were performed in accordance with the guidelines of the French Agriculture and Forestry Ministry for handling animals (APAFIS #26 770) and the official European guidelines (Directive 2010/63/UE). Male and female D1-GFP mice (Tg(Drd1-EGFP)X60Gsat) were used, aged between P16 and P25 for young mice and between 6 to 10 weeks for adult mice. Young mice from P16 to P25 have been chosen, as the number of layer V D1 receptor sites is at its maximum at these stages of development in the prefrontal cortex (Leslie et al., 1991). D1-GFP mice express the GFP under the D1 receptor promoter, enabling the identification of D1-expressing cells. Mice were housed collectively under artificial conditions of light (12/12 h light/dark cycle, light on at 7:00 a.m.), with food and water access *ad libitum*.

### Slice preparation

Mice were deeply anesthetized using ketamine and xylazine (100 and 20 mg/kg, i.p., respectively). After the disappearance of all arousal reflexes, a thoracotomy was done to enable the transcardial perfusion of an ice-cooled and oxygenated with carbogen (95% O_2_/5% CO_2_) cutting solution containing 250 mM sucrose, 2.5 mM KCl, 1.25 mM NaH2PO4·H2O, 0.5 mM CaCl2·H2O, 10 mM MgSO4·7H2O, 10 mM D-glucose and 26mM NaHCO3. The brain was then quickly removed and glued to the stage of a vibratome (VT1200S; Leica Microsystems, Germany) and placed into a cutting chamber filled with the cutting solution and oxygenated with carbogen. The brain was then cut into 300 µm thick sections, which were then incubated for 1 hour into a 37°C warmed ACSF containing 126 mM NaCl, 2.5 mM KCl, 1.25 mM NaH2PO4·H2O, 2 mM CaCl2·H2O, 2 mM MgSO4·7H2O, 26 mM NaHCO3, and 10 mM D-glucose, 1 mM sodium pyruvate and 4.9 µM L-glutathione reduced and oxygenated with carbogen. Slices were then placed at room temperature for 30 minutes before recordings.

### Drugs

Drugs were prepared in double-distilled water as concentrated stock solutions, then aliquoted and stored at -20°C. Drugs were diluted daily at the experimental concentrations and perfused in the recording chamber. In all experiments, glutamatergic AMPA/kainate and NMDA receptors were blocked with 20 µM 6,7-dinitroquinoxaline-2,3-dione (DNQX, Tocris, UK) and 50 µM D-(-)-2-amino-5-phosphonopentanoic acid (AP V, Tocris, UK) respectively, and GABA_A_ receptors were blocked using 10 µM 6-Imino-3-(4-methoxyphenyl)-1(6*H*)-pyridazine butanoic acid hydrobromide (GABAzine, Tocris, UK). To block D1 receptors, 1 µM D1 receptor antagonist (*R*)-(+)-7-Chloro-8-hydroxy-3-methyl-1-phenyl-2,3,4,5-tetrahydro-1*H*-3-benzazepine hydrochloride (SCH 23390, Sigma, France) was used, and to activate D1 receptors, 2.5 µM D1 receptor agonist (±)-6-Chloro-2,3,4,5-tetrahydro-1-phenyl-1*H*-3-benzazepine hydrobromide (SKF 81297, Tocris, UK) was used. Electrophysiological recordings were made 20 minutes after drug application.

### *Ex vivo* electrophysiological recordings

Single slices were placed in a recording chamber continuously perfused with a recording solution containing 126 mM NaCl, 3 mM KCl, 1.25 mM NaH2PO4·H2O, 1.6 mM CaCl2·H2O, 2 mM MgSO4·7H2O, 10 mM D-glucose and 26 mM NaHCO3, oxygenated with carbogen and heated at 32°C. PNs were visualized under IR-DIC and fluorescence microscopy using a 63X water-immersion objective (W Plan-Apochromat 63X/1.0 VIS-IR, Zeiss) equipped on an axio examiner Z.1 microscope (Zeiss, Germany). PNs were identified by the shape of their cell bodies and then confirmed by their electrophysiological signature. Recordings of PNs were made using patch pipettes of impedance between 4-9 MΩ. These pipettes were made from glass capillaries (GC150F10; Warner Instruments, Hamden, CT, USA) pulled with a horizontal pipette puller (P-97; Sutter Instruments, Novato, CA, USA). All recordings were made in the whole-cell configuration using an internal pipette solution containing 135 mM K-gluconate, 3.8 mM NaCl, 1 mM MgCl2·6H2O, 10 mM HEPES, 0.1 mM Na4EGTA, 0.4 mM Na2GTP, 2 mM MgATP and 5.4 mM biocytin (pH = 7.2, ∼292 mOsm). Recordings were corrected for a junction potential of -13 mV. Experiments were done with a Multiclamp 700B amplifier and digidata 1550B digitizer controlled by clampex 11.0 (Molecular Devices LLC). Recordings were acquired at 20 kHz and low-pass filtered at 4 kHz. Series resistance was monitored throughout the experiment by voltage steps of -5 mV, and data were discarded when the series resistance changed by > 20%. After recordings, slices were fixed for 24 hours at 4°C in a solution of PBS 0.01 M containing 4% of paraformaldehyde, then washed and stored in PBS 0.01 M containing 0.03% of sodium azide until further histological processing.

### Histology

Transcardial perfusions were made on mice following the same procedure as described for slice preparation, except the ACSF used did not contain sodium pyruvate and glutathione. The brains were then post-fixed at 4°C in a solution of PBS 0.01 M containing 4% paraformaldehyde for 24 hours, washed, and cut into 50 µm thick slices with a vibratome (VT1000S; Leica Microsystems, Mannheim, Germany). Slices were then processed for immunohistochemistry labeling. Slices were placed in a blocking buffer for 2 hours, then 48 hours in a solution of PBS 0.01M/Triton X-100 0.3% containing the primary antibodies (Table 1). Slices were washed three times in PBS 0.01 M, incubated with the secondary antibodies for 2 hours, washed three times again with PBS 0.01 M, and then mounted onto slides in DAPI fluoromount medium (SouthernBiotech). Images were taken with a confocal microscope (Leica TCS SP8, Leica Microsystems, Mannheim, Germany) equipped with an HC PL APO 20x/0.75 IMM CORR CS2 objective (used to take pictures for counting) and an HC Plan Apo CS2 63X oil NA 1.40 objective (used to take patched neurons pictures). Confocal images were further processed using Fiji. Counting and colocalization were made manually using a rectangle-delimited M1 area. Layers were delimited by the DAPI and Ctip2 labeling. The delimitation between layer I and layer II-III was placed where a sharp decrease in nuclear DAPI labeling is found. The layer V was placed where there is an increase in Ctip2 labeling intensity. Three slices containing a large part of M1 from three mice were used for counting (Fig. 1 A,B, 2 A, B).

**Figure 1:**
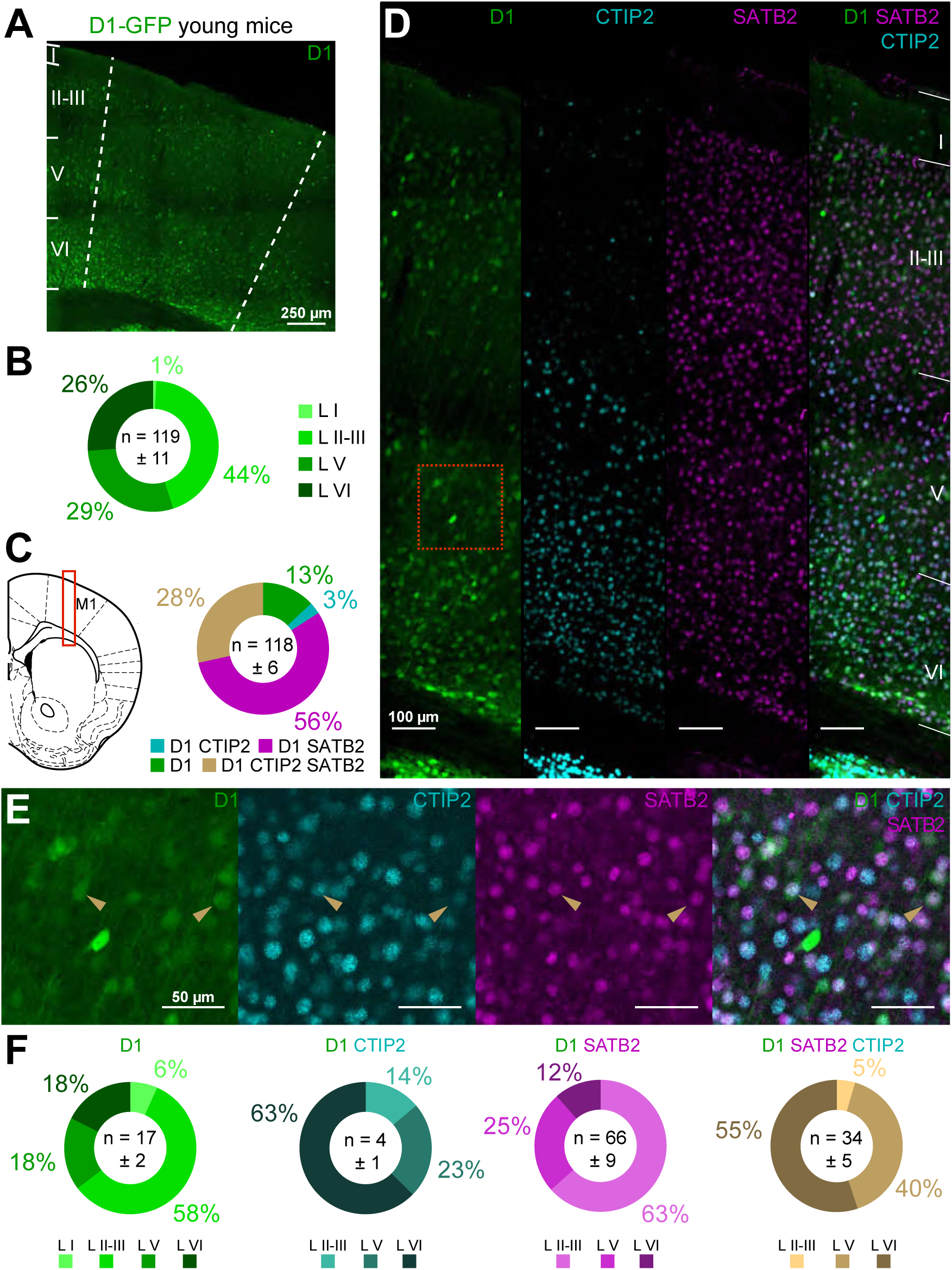
Distribution of D1 receptor-expressing neurons in M1 of young mice. **A.** Image of a coronal section at the level of M1 showing the D1-expressing neurons in a young animal. **B.** Laminar distribution of D1-positive cells in M1 of young mice. For each category, the darker the color, the deeper the layer. **C.** Left, schematic of a coronal slice containing M1. The red rectangle indicates the area imaged in D. Right, distribution in % of all D1-expressing cells in M1 according to their molecular identity. **D.** Example of the labeling obtained for D1 (green), Ctip2 (blue), and Satb2 (magenta) in M1. **E.** Enlarged view of layer V at the level of the red-dotted square in D for each molecular marker. The brown arrowheads indicate neurons positive for D1, Ctip2 and Satb2 labeling. **F.** Distribution of D1 positive only (green), D1 and Ctip2 positive only (blue), D1 and Satb2 positive only (magenta), and D1, Ctip2 and Satb2 positive (brown) cells in M1 layers. For each category, the darker the color, the deeper the layer. Data are given as mean ± SEM.

**Figure 2:**
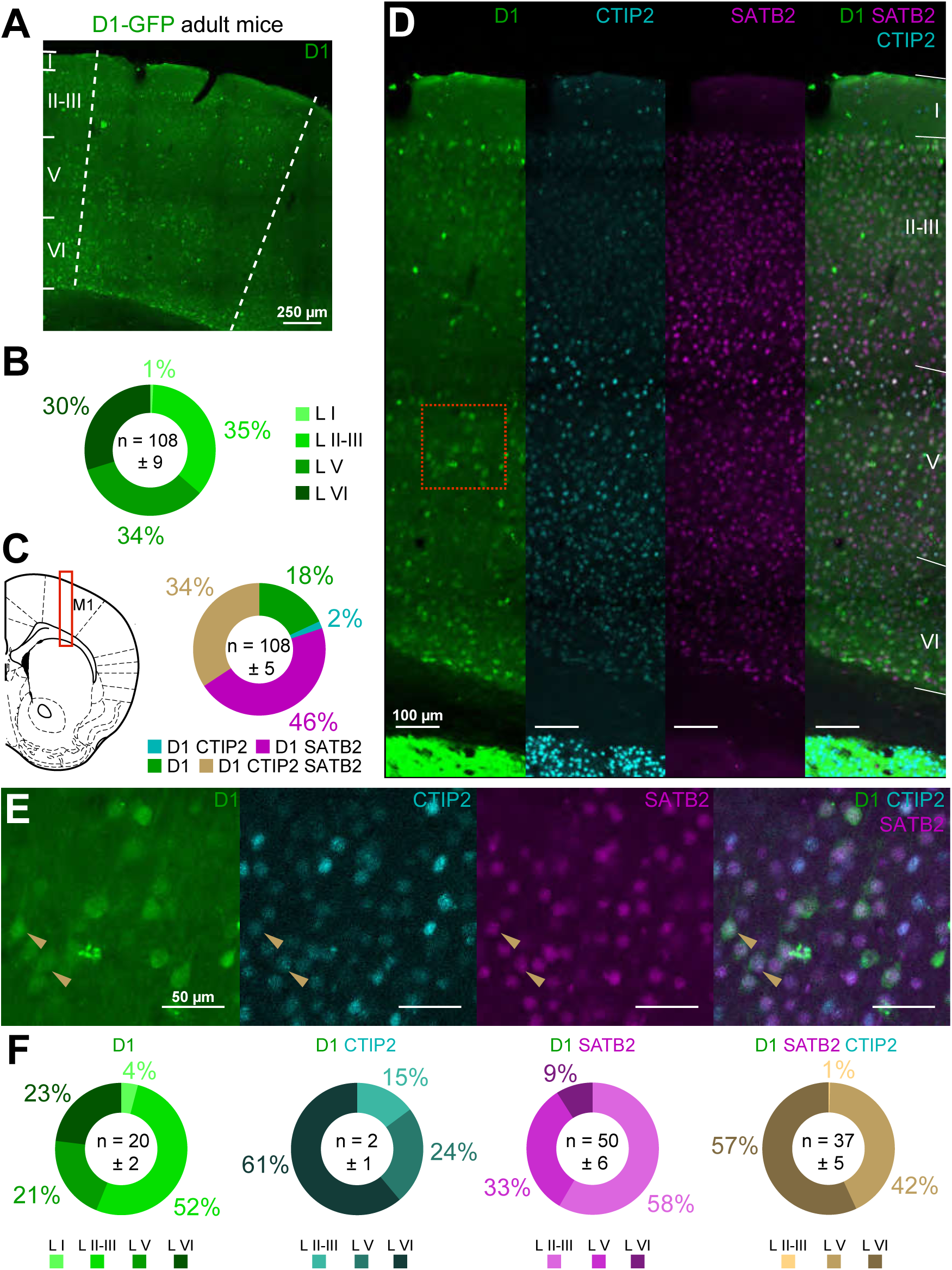
Distribution of D1 receptor-expressing neurons in M1 of adult mice. **A.** Image of a coronal section at the level of M1 showing the D1-expressing neurons in the brain of an adult mouse. **B.** Laminar distribution of D1-positive cells in M1 of adult mice. For each category, the darker the color, the deeper the layer. **C.** Left, schematic of a coronal slice containing M1. The pictures in D. are taken from the area delineated by the red-dotted square. Left, distribution in % of all D1-expressing cells in M1 according to their molecular identity in adult mice. **D.** Example of the labeling obtained for D1 (green), Ctip2 (blue), and Satb2 (magenta) in M1 of an adult mouse. **E.** Higher magnification of the layer V of M1 at the level of the red-dotted square in B. for the same molecular markers. The brown arrowheads indicate neurons positive for D1, Ctip2 and Satb2 labeling. **F.** Distribution of D1 positive only (green), D1 and Ctip2 positive only (blue), D1 and Satb2 positive only (magenta), and D1, Ctip2 and Satb2 positive (brown) cells in M1 layers. For each category, the darker the color, the deeper the layer. Data are given as mean ± SEM.

**Table 1:**
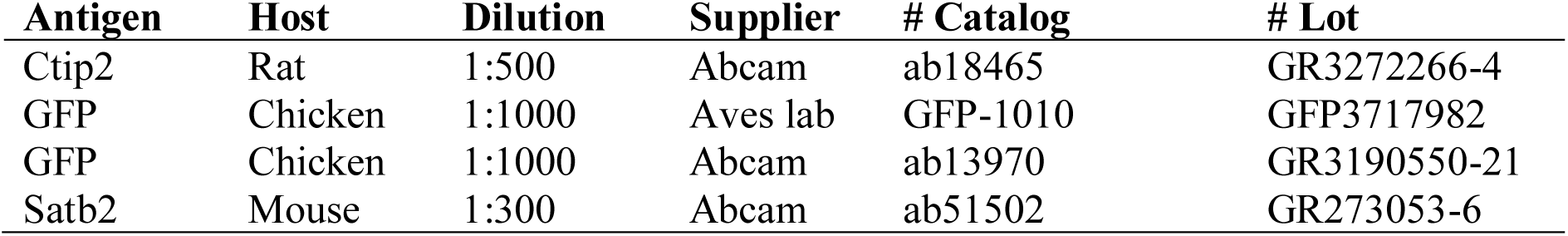
List of primary antibodies used

### Data analysis

Electrophysiological data were analyzed using Clampfit 10.7 (Molecular Devices, USA) and Origin 7 (OriginLab, USA). Input-output (F-I) curves were generated by injecting increasing 1s depolarizing currents (25 pA increments, from -150 pA to 225 pA) and counting the number of evoked action potentials. Input resistance was calculated using Ohm’s law when a current of -50 pA was injected. Δ*U* corresponds to the voltage variation between the baseline and the new voltage recorded due to the current injection. The rheobase was determined by injecting increasing depolarizing currents of 500 ms, with 1 pA increments. Action potential half width and peak amplitude were obtained after detecting each spike with the threshold search in Clampfit 10.7. The action potential threshold was measured as the beginning of the rising slope of the phase plots of the neurons. These phase plots were made at rheobase using Clampfit 10.7. Electrophysiological traces were processed using Origin 7.

### Statistics

Statistical analyses were performed using Prism 9.3.1 (GraphPad Software Inc). For paired analysis (*i.e.*, for membrane resting potential, rheobase, action potential peak amplitude, action potential threshold, input resistance, and action potential half-width) Wilcoxon signed rank tests (WSR) were performed. In this case, the black dotted line represents the mean ± standard error to the mean (SEM) of all neurons, and the transparent-colored lines represent individual neurons. For firing frequency, two-way multiple comparisons ANOVA followed by a Bonferroni post hoc were made. In all tests, the level of significance was set at p < 0.05. Details about statistical tests and p-values are shown in Table 2. Data are represented as mean ± SEM.

**Table 2:**
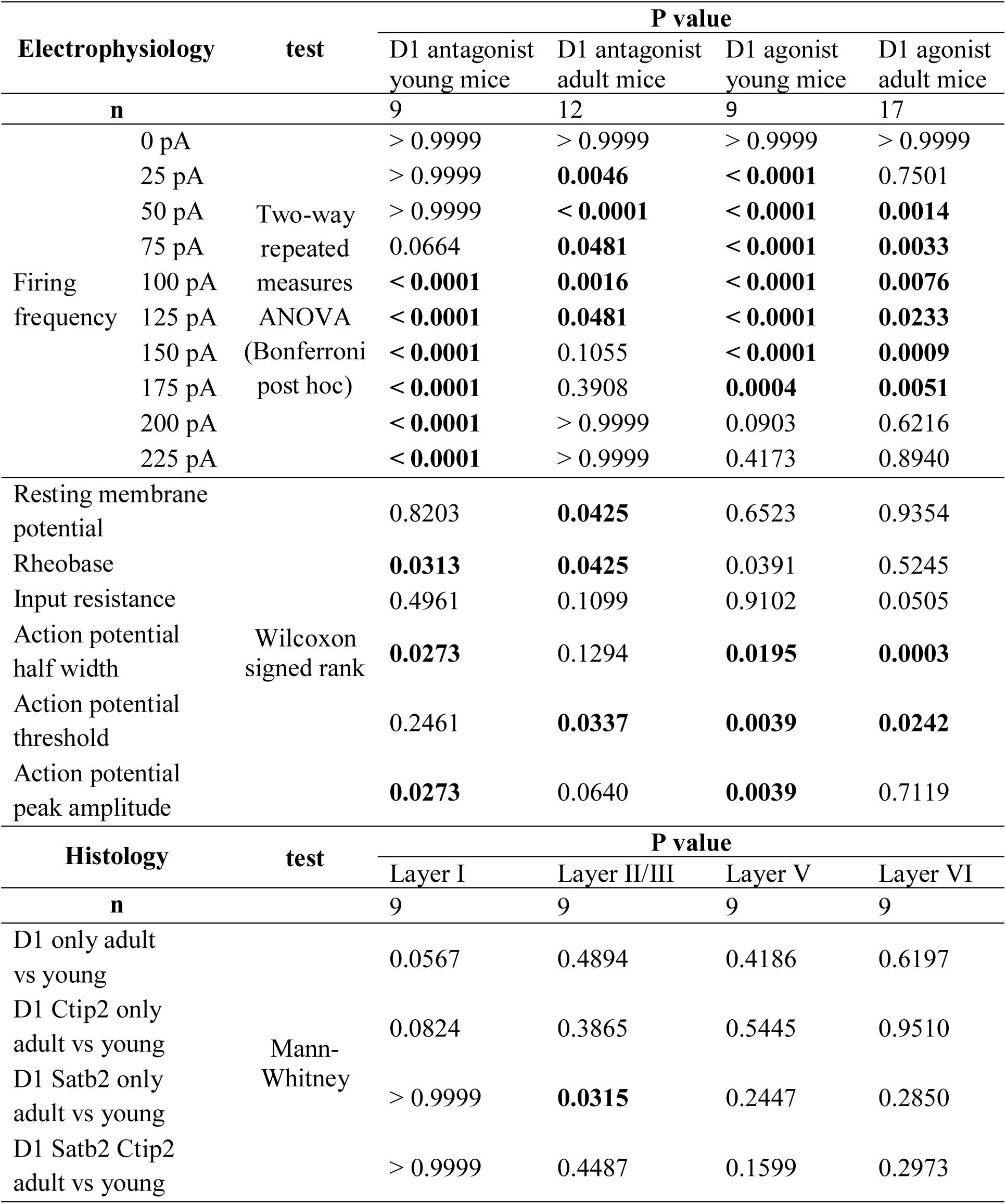
Summary of the statistics of the study

## Results

### The distribution of D1 receptors expressing cells in M1 is similar in young and adult mice

Taking advantage of the D1-GFP transgenic mice in which the GFP is expressed under the control of the D1 receptor promoter, we first performed a quantitative layer-based mapping of M1 neuronal populations expressing the D1 receptor in young and adult mice. The analysis of the GFP fluorescence in M1 brain slices revealed that D1 receptor-expressing cells were distributed in all M1 cortical layers and with a similar distribution in young and adult mice (Fig. 1A, B; 2A, B). In young mice, ∼26% of D1 receptor-expressing cells were localized in layer VI, ∼29% in layer V, 44% in layers II/III, and less than 1% in layer I (Fig. 1A, B). In adult mice, ∼30% of cells expressing the D1 receptor were localized in layer VI, ∼34% in layer V, ∼36% in layers II/III and less than 1% in layer I (Fig. 2A, B).

To refine the molecular identity of the D1 receptor-expressing cells, we performed immunostaining to quantify the colocalization of GFP with specific markers of two classes of pyramidal neurons. Ctip2 and Satb2 transcription factors were used as molecular markers of PT and IT neurons, respectively (Arlotta et al., 2005; Molnár and Cheung, 2006; Alcamo et al., 2008; Britanova et al., 2008; Digilio et al., 2015). The D1 receptor-expressing neurons were then divided into four categories: the cells expressing only the D1 receptor, neurons expressing the D1 receptor and only Ctip2, those expressing the D1 receptor and only Satb2, and those expressing the D1 receptor and both Ctip2 and Satb2, in young and adult mice (Fig. 1C-F; 2C-F). Most of the cells expressing the D1 receptor in M1 co-expressed Satb2 (around 80% both in young and adult mice) and very few cells co-expressed only Ctip2 (3.3% and 1.83% in young and adult mice, respectively) (Fig. 1C; 2C). The laminar distribution of the cells in the four categories was also similar in young and adult mice (Fig. 1D-F; 2D-F). The cells expressing only the D1 receptor were mostly localized in layer II-III, 58.01% in young (Fig 1D-F) and 52.09% in adult mice (Fig. 2D-F). The few D1 receptor-positive cells co-expressing only Ctip2 were mainly localized in layer VI both in young (Fig. 1D-F) and adult mice (Fig. 2D-F). The D1 receptor-positive cell co-expressing Satb2 were mainly localized in layer II-III as they represented 63.46% in young (Fig 1D-F) and 58.68% in adult mice (Fig. 2D-F) and in layer V (25.03% and 32.63%). Finally, a non-negligible number of D1 receptor positive-cells co-expressing Ctip2 and Satb2 were also counted. They were mainly found in layer V of young (55.15%) and adult (56.86%) mice (Fig. 1D-F; 2D-F).

### D1 receptor activation increases PNs excitability both in young and adult animals

We then explored if the DA D1 receptor can modulate the intrinsic electrical properties of individual neurons and whether DA modulation of these neurons changes with age. We focused our attention on PNs in layer V, the main output layer of the cortex (Lévesque et al., 1996; Veinante et al., 2000; Hattox and Nelson, 2007; Aronoff et al., 2010, reviewed in Harris and Shepherd, 2015) which is largely innervated by DA fibers (Vitrac et al., 2014). Using patch-clamp recording, we first investigated *ex vivo* the effects of the activation of the D1 receptors on PNs’ intrinsic electrical properties in M1 layer V (Fig. 3, 4). PNs were identified on morphological (triangle shape of their cell bodies) and electrophysiological criteria. To prevent a network effect, intrinsic properties of PNs were recorded while pharmacologically blocking fast glutamatergic and GABAergic transmission using DNQX (50 µM), APV (20 µM), and GABAzine (10 µM). In young animals (Fig 3), bath application of the D1 agonist SKF 81297 (2.5 µM) changed the intrinsic properties of the PNs recorded. Many parameters were measured. We observed that PNs were more excitable as they fired more action potentials in response to current injections from 25 pA to 175 pA (Fig. 3B, two-way repeated measures ANOVA, F_(9, 72)_ = 138.5, p < 0.0001, n = 9) in presence of the D1 receptor agonist SKF 81297 (Fig. 3C). Furthermore, the rheobase, action potential threshold, half-width and peak amplitude were significantly lower with the application of D1 receptor agonist SKF 81297 compared to control conditions (Fig. 3D, Wilcoxon signed rank (WSR), p < 0.05, n = 9). No significant effect was observed concerning the resting membrane potential and the input resistance (Fig. 3D, WSR, p > 0.05, n = 9).

**Figure 3:**
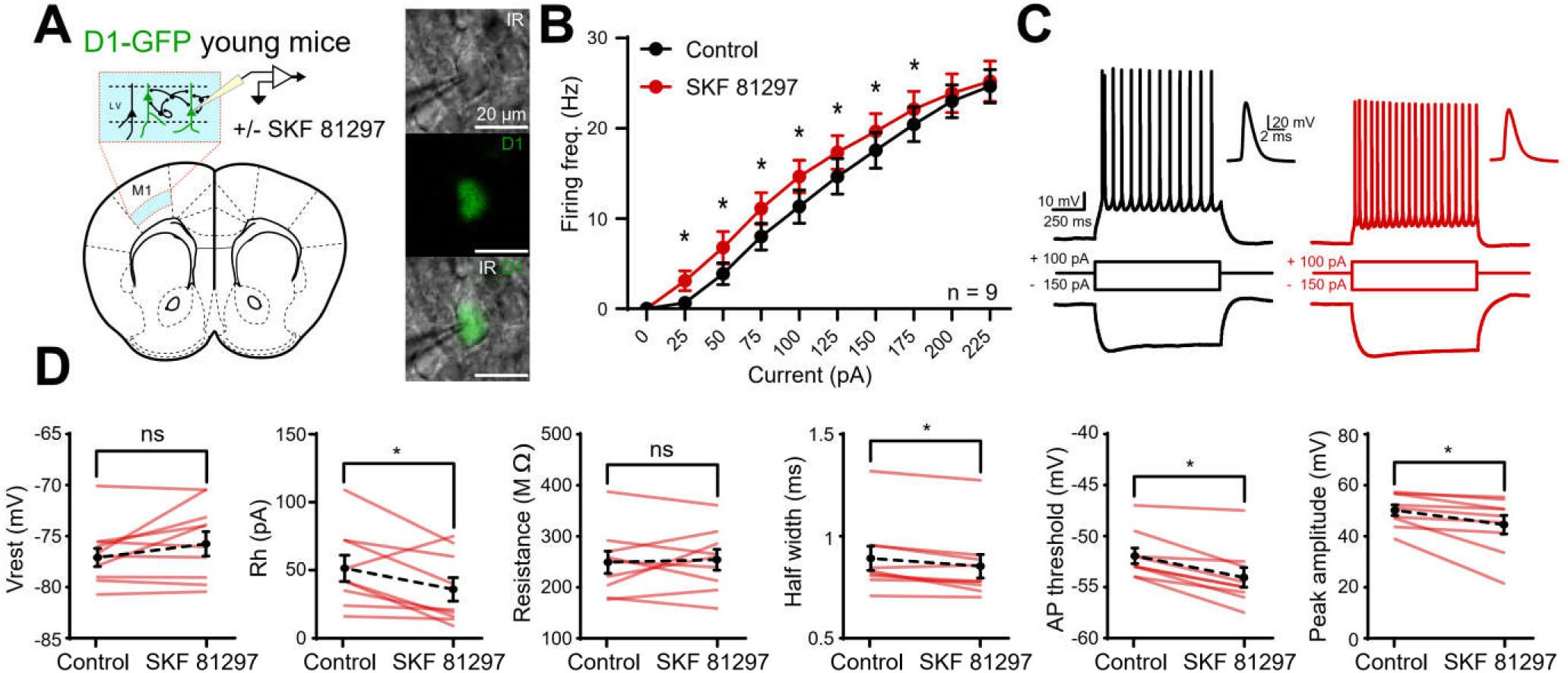
Effect of the D1 dopaminergic agonist SKF 81297 on layer V M1 pyramidal cells’ excitability in young mice. **A.** Left, experimental design. Right, images of a recorded pyramidal neuron expressing the D1 receptor under IR-DIC (top, IR), fluorescence (GFP, middle) and merge of the two pictures (down, IR/GFP). **B.** Input/output curves in control (black) and in presence of the D1 agonist (red). n = 9. * p < 0.05 (two-way repeated measures ANOVA). **C.** Responses to depolarizing and hyperpolarizing current steps in an individual pyramidal neuron recorded before (left) and after bath application of D1 agonist (right, in red). An expanded view of a single spike is presented next to each trace. **D.** Cell parameters recorded before and after bath application of D1 agonist, from left to right: resting membrane potential, rheobase, input resistance, half-width of action potentials, action potential threshold, and peak amplitude of action potential. n = 9. * p < 0.05, ns = non-significant (WSR).

The effect of D1 receptor activation on layer V M1 PNs was then assessed in adult mice (Fig. 4). As for young mice, D1 agonist increased the excitability of PNs as illustrated by the recording of a PN in response to a 100 pA current injection (Fig. 4C). For current injections from 50 pA to 175 pA, PNs fired more action potentials in presence of SKF 81297 compared to control conditions (Fig. 4B, two-way repeated measures ANOVA, F_(9, 144)_ = 306.2, p < 0.0001, n = 17). Moreover, the input resistance, the action potential half width, and the action potential threshold were significantly lower in the presence of SKF 81297 compared to the control condition (Fig. 4D, WSR, p < 0.05, n = 17). No significant effect was observed concerning the resting membrane potential, the rheobase, and the action potential peak amplitude (Fig. 4D, WSR, p > 0.05, n = 17). These effects were specific to the activation of the receptor as coactivating and blocking the D1 receptor simultaneously did not affect significantly the intrinsic properties of the PNs (Supplementary Fig 1).

**Figure 4:**
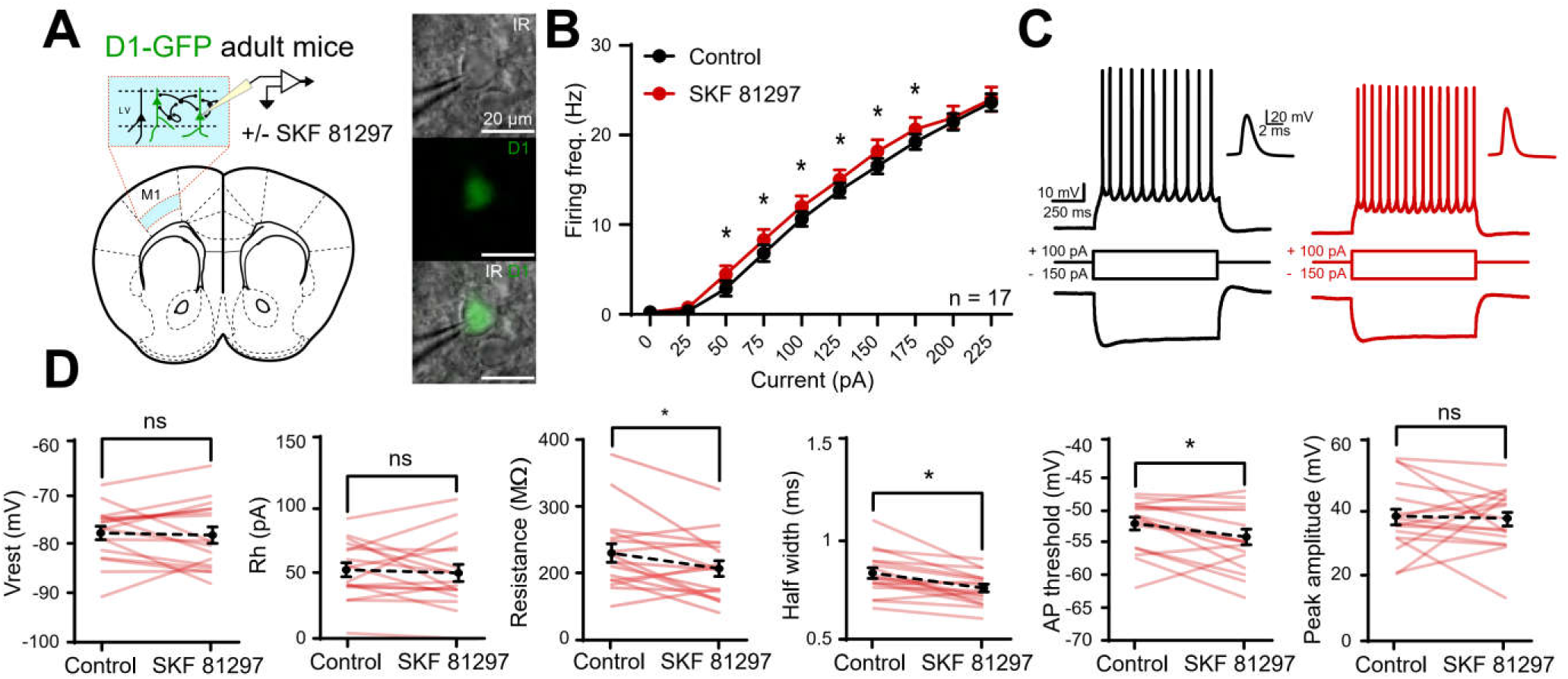
Effect of D1 dopaminergic agonist SKF 81297 on layer V M1 pyramidal cells’ excitability in adult mice. **A.** Left, experimental design. Right, images of a recorded pyramidal neuron expressing the D1 receptor under IR-DIC (top), fluorescence (middle, GFP) and merge of the two pictures (down). **B.** Input/output curve in control (black) and in presence of the D1 agonist (red). n = 17. * p < 0.05 (two-way repeated measures ANOVA). **C.** Responses to depolarizing and hyperpolarizing current steps in an individual pyramidal neuron recorded before (left) and after bath application of D1 agonist (right). An expanded view of a single spike is presented next to each trace. **D.** Cell parameters recorded in adult animals before and after bath application of D1 agonist, from left to right: resting membrane potential, rheobase, input resistance, half-width of action potentials, action potential threshold, and peak amplitude of action potential. n = 17. * p < 0.05, ns = non-significant (WSR).

### Blockade of D1 receptor differently impact layer V PNs intrinsic properties according to the age

We then investigated the effect of blocking the D1 receptor on PNs’ intrinsic properties in M1 layer V (Fig. 5 and 6). In young animals (Fig 5), bath application of the D1 antagonist SCH 23390 (1µM) had the opposite effect of the bath application of the D1 agonist on the intrinsic properties of the PNs recorded. Indeed, we observed a significant decrease in the neuronal excitability of PNs in the presence of SCH 23390. PNs fired fewer action potentials in response to a somatic injection of depolarizing currents in the presence of SCH 23390 compared to control conditions (Fig. 5B, two-way repeated measures ANOVA, F_(9, 72)_ = 48.58, p < 0.0001, n = 9), as illustrated by the recorded traces (Fig. 5C) and by the frequency/current input-output curve (Fig. 5B). Furthermore, the rheobase of these neurons was significantly higher with SCH 23390 compared to control conditions (WSR, p < 0.05, n = 9), and the input resistance, the action potential half-width and peak amplitude were significantly lower with SCH 23390 compared to control conditions (Fig. 5D, WSR, p < 0.05, n = 9). No significant differences were observed concerning the resting membrane potential and the action potential threshold between SCH 23390 and control conditions (Fig. 5D, WSR, p > 0.05, n = 9).

**Figure 5:**
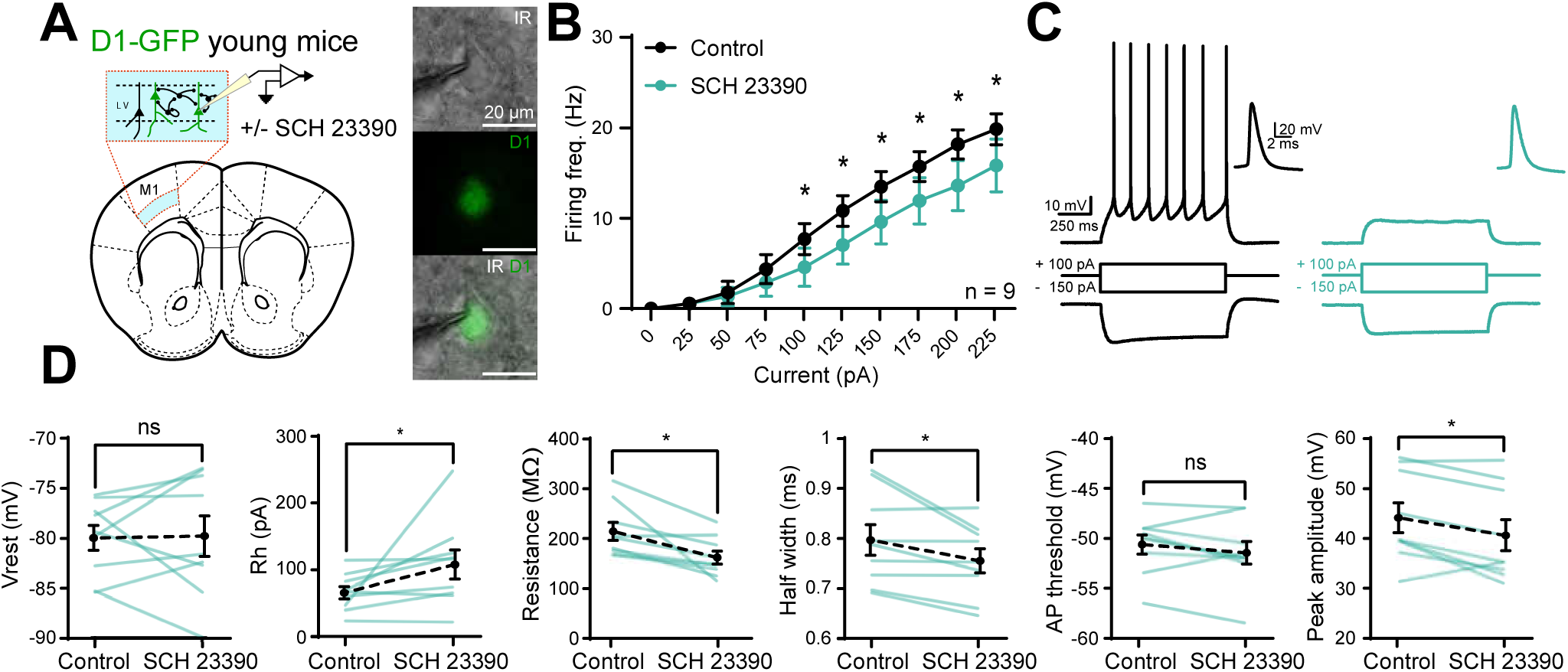
Effect of D1 dopaminergic antagonist SCH 23390 on layer 5 M1 pyramidal cells’ excitability in young mice. **A.** Left, experimental design. Right, images of a recorded pyramidal neuron expressing the D1 receptor under IR-DIC (top), fluorescence (middle) and merge of the two pictures (down). **B.** Input/output curves in control (black) and in presence of the D1 antagonist (blue). n = 9. * p < 0.05 (two-way repeated measures ANOVA). **C.** Responses to depolarizing and hyperpolarizing current steps in an individual pyramidal neuron recorded before (left) and after bath application of D1 antagonist (right in blue). An expanded view of a single spike is presented next to each trace. **D.** Cell parameters recorded in young animals before and after bath application of D1 antagonist, from left to right: resting membrane potential, rheobase, input resistance, half-width of action potentials, action potential threshold and peak amplitude of action potentials. n = 9. * p < 0.05, ns = non-significant (WSR).

**Figure 6:**
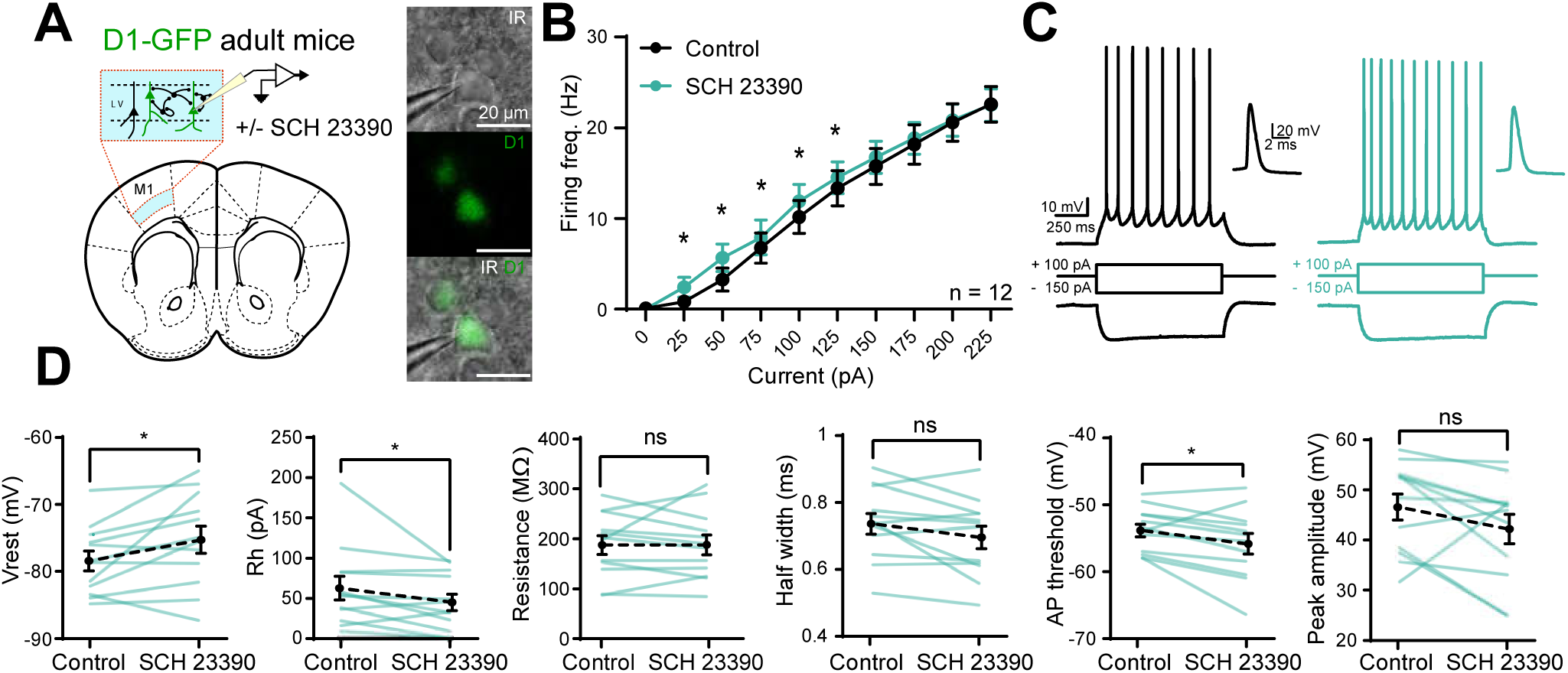
Effect of D1 dopaminergic antagonist SCH 23390 on layer 5 M1 pyramidal cells’ excitability in adult mice. **A**. Left, experimental design. Right, images of a recorded pyramidal neuron expressing the D1 receptor under IR-DIC (top), fluorescence (middle) and merge of the two pictures (down). **B.** Input/output curves in control (black) and in presence of the D1 antagonist (blue). n = 12. * p < 0.05 (two-way repeated measures ANOVA). **C.** Responses to depolarizing and hyperpolarizing current steps in an individual pyramidal neuron recorded before (left) and after bath application of D1 antagonist (right in blue). An expanded view of a single spike is presented next to each trace. **D.** Cell parameters recorded in adult animals before and after bath application of D1 antagonist, from left to right: resting membrane potential, rheobase, input resistance, half width of action potential, action potential threshold and peak amplitude of action potential. n = 12. * p < 0.05, ns = non-significant (WSR).

The same experiments were then performed in adult mice (Fig. 6). Surprisingly, the excitability of layer V PNs was increased by the bath application of the D1 receptor antagonist as it was with the application of the D1 receptor agonist. Indeed, the recorded PNs fired more action potentials following low-intensity stimulation ranging from 25 pA to 125 pA with 1 µM SCH 23390 than in control conditions (Fig. 6B, C, two-way repeated measures ANOVA, F_(9, 99)_ = 124.4, p < 0.0001, n = 12). Moreover, the resting potential of these neurons was more depolarized in the presence of D1 receptor antagonist SCH 23390 compared to control conditions (Fig. 6D, WSR, p < 0.05, n = 12). Furthermore, the rheobase and the action potential threshold of layer V M1 PNs were lowered while blocking D1 receptors (Fig. 6D, WSR, p < 0.05, n = 12). No significant effects were observed concerning the input resistance, the action potential half-width, and peak amplitude (Fig. 6D, WSR, p > 0.05, n = 12).

## Discussion

DA signaling is crucial for the control of voluntary movement and for motor learning, however, how D1 receptors modulate intrinsic properties of individual neurons in mouse primary motor cortex was poorly understood and contradictory. In this study, we demonstrated *ex vivo* that neurons expressing the D1 receptor are widely distributed in all layers of M1 similarly in young and adult mice. Moreover, we showed that blocking or activating the D1 receptor modulates in a specific way the intrinsic properties of layer V PNs depending on the age of the animals.

We first showed that D1 receptor-expressing cells are widely distributed in all layers of M1. This distribution correlates well with the localization of the DAergic fibers in the superficial and deep layers of M1 (Berger et al., 1985; Descarries et al., 1987; Raghanti et al., 2008; Vitrac et al., 2014). As it has been shown in the medial prefrontal cortex that the expression of the D1 receptor changes during postnatal development (Leslie et al., 1991), we looked at the expression of the D1 receptor in M1 at two stages. The mapping of these D1 receptor-expressing cells reveals a similar distribution regardless of the age of the mice, with however a slight tendency to decrease in superficial layers and to increase in deep layers when mice get older. Projection neurons progressively acquire subtype and area identities by transcriptional mechanisms (Greig et al., 2013). To better characterize the identity of the neurons that express the D1 receptor, we used the classical biological markers of distinct pyramidal neurons, Ctip2 for PT neurons and Satb2 for IT neurons (Arlotta et al., 2005; Molnár and Cheung, 2006; Alcamo et al., 2008; Britanova et al., 2008; Digilio et al., 2015). Satb2 represses the expression or prevents the activity of Ctip2 (Alcamo et al., 2008; Britanova et al., 2008). Thus, overexpression of Satb2 during adolescence in layer II/III could be of importance to repress the expression of other transcriptional factors leading to the specification of neurons other than subcortical- and callosal-projection neurons. Nearly 15% of the cells were expressing only the D1 receptor, suggesting that some inhibitory interneurons and some CT pyramidal neurons in layer VI also express the D1 receptor. As very few D1 receptor-expressing neurons express only Ctip2, it indicates that the majority of PT neurons do not have the D1 receptor as already reported (Gaspar et al., 1995; Shepherd, 2013). In any case, most of the D1-expressing cells also express Satb2 suggesting that a majority of these cells are IT neurons. Because the D1/Satb2 cells are located in different layers, they could be further identified as IT Cortico-cortical neurons in layer II/III and IT Cortico-striatal neurons in deeper layers (Huang et al., 2013; Shepherd, 2013). As it has already been reported in neocortical regions in mice (McKenna et al., 2011), staining with Ctip2 antibodies revealed neurons that expressed high levels of Ctip2 protein while Ctip2 expression level was much lower in others. Satb2 is known to negatively regulate the level and activity of Ctip2 in neurons (Alcamo et al., 2008; Britanova et al., 2008). Interestingly, some cells co-express Satb2 and Ctip2 in deep layers highlighting the existence of a subpopulation of neurons that have been already described in the somatosensory cortex (Harb et al., 2016), motor area (Sohur et al., 2014; Tantirigama et al., 2014) and hippocampus (Lickiss et al., 2012; Nielsen et al., 2014; Digilio et al., 2015) that also express the D1 receptor.

Using a combination of pharmacology and *ex vivo* electrophysiology, we studied how DA modulates the intrinsic properties of PNs in layer V of M1 in young and adult mice. Intracellular cascades induced by DAergic receptor activation vary with the cell types and the brain region (Stoof and Kebabian, 1981; Sidhu et al., 1991; Rioult-Pedotti et al., 2015; Mishra et al., 2018) and more importantly, the DA receptor can be coupled with several G proteins (Sidhu, 1998). To avoid a network effect and to specially study the impact of the D1 receptor on the electrical intrinsic properties of the neurons, we isolated the neurons recorded from the network by the presence of fast synaptic transmission blockers. In this study, we demonstrated that activating the D1 receptor increased the excitability of M1 layer V PNs, presumably IT, both in young and adult mice. However, even if we observed a significant global change in intrinsic properties, we observed an important inter-individual variability of the responses induced by the bath application of the agonist. Even if the immunohistochemistry experiments (Fig. 1, 2) indicate that most of the D1 receptor-expressing cells in layer V of M1 are IT neurons, this variability suggests that subtypes of PNs expressing D1R have been recorded (Sohur et al., 2014; Tantirigama et al., 2014). At first glance, these results do not seem to agree with an *in vivo* study showing a decrease in excitability of layer V PNs following DA local application in rat motor cortex (Awenowicz and Porter, 2002), but this study was targeting the PT neurons.

Interestingly, we demonstrated an age-dependent action of D1 receptor antagonist on intrinsic electrical properties of layer V PNs. While D1 receptor blockade decreased the excitability of M1 layer V PNs in young animals, it increased their excitability in adults. In young mice, as the D1 receptor activation induced an increase in M1 layer V PNs excitability, it was consistent to observe a decrease in the excitability of M1 layer V PNs by the blockade of the D1 receptor. This supports the idea that the D1 receptors recruit an excitatory G protein. Surprisingly, in adults, the D1 receptor blockade increased the firing frequency and lowered the action potential threshold of M1 layer V PNs as it was for the activation of the D1 receptor. Even if it is surprising, the effect observed in adults is in line with the work of Swanson and colleagues who recently showed *ex vivo* that D1 receptor antagonism caused increased excitability of layer V PNs (Swanson et al., 2021). This electrophysiological signature in the presence of the antagonist reminds the altered electrophysiological properties (*i*.*e*., higher firing frequency and depolarized resting membrane potential) described *in vivo* in cortical neurons of Parkinsonian rats in which the DAergic transmission was interrupted (Degos et al., 2013). Besides, drug effects on receptor activity are often integrated with some preexisting level of receptor activity which depends on endogenous ligand and constitutive receptor activity. It has been already reported that the D1 receptor in other brain regions can have a constitutive activity (Rankin et al., 2006; Zhang et al., 2014) meaning a receptor activity in the absence of ligands at the binding site. The opposite effect observed in young and adult mice may suggest that the D1 receptor can be constitutively active in adults but not in young animals. This hypothesis is also reinforced by the fact that the bath application of the antagonist of the D1 receptor in adults produces the same effect as the bath application of the agonist. Besides, the effects observed in adults by the bath application of D1 receptor agonist were not as strong as the ones observed in young animals. If the constitutive activity of the receptor in adult is high, further increasing the activity of the receptor by the agonist may have a small effect relative to the baseline and may be harder to detect. Additionally, the effect of the antagonist will work if high level of DA, the data obtained can also suggest a different DAergic tone in young and adult mice that could arise from different emotional or motor states. Thus, the DA modulation may be governed by different mechanisms at different ages

In summary, this study unravels the impact of D1 receptors on M1 layer V PNs *ex vivo*, and maps for the first time the D1 receptor-expressing neurons in M1 according to their molecular profile. The D1 receptors modulation of M1 layer V PNs is of importance for the physiological completion of M1 processes, such as motor learning and execution of fine motor tasks in a healthy M1 and impaired DA signaling will lead to pathologies.

## Abbreviations

DA: dopamine
DAergic: dopaminergic
GFP: green fluorescent protein
M1: primary motor cortex
PNs: pyramidal neurons

## Acknowledgments

This work was supported by the University of Bordeaux, the CNRS (Centre National de la Recherche Scientifique), the French government through the University of Bordeaux’s IdEx "Investments for the Future" program/GPR BRAIN_2030 (to J.B. and M. L. B.J.), and the Bordeaux Neurocampus Department (Seed project Damoco). V.P. benefitted from the help of the Bordeaux Neurocampus Graduate Program, managed by the French National Research Agency reference ANR-17-EURE-0028. Confocal images were taken at the Bordeaux imaging center.

## Figure legends

**Supplementary figure 1:**
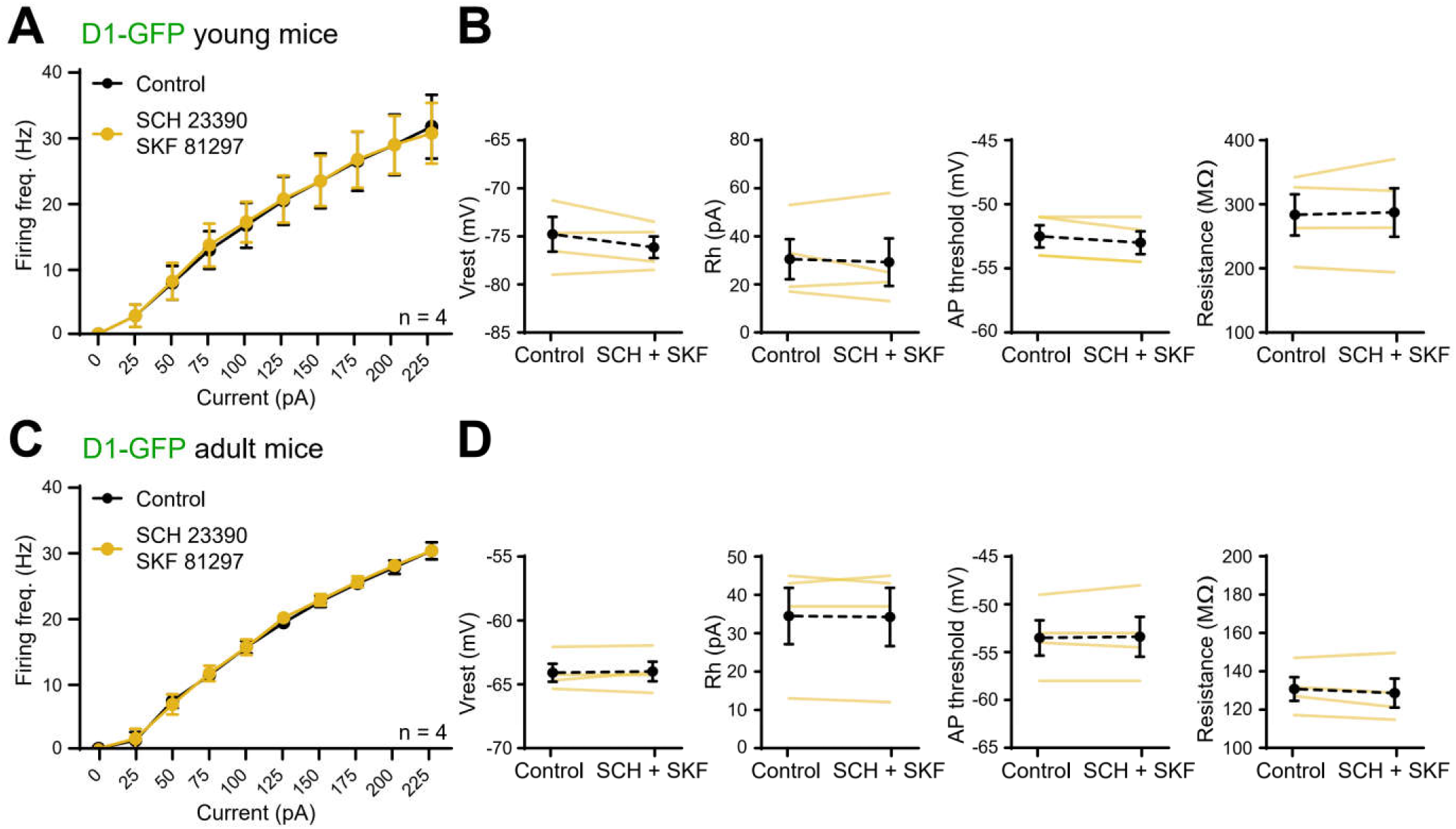
Effect of the co-application of D1 agonist SKF 81297 and D1 antagonist SCH 23390 on layer 5 M1 pyramidal cells’ excitability. **A.** Input/output curves in control and in the presence of both D1 agonist and antagonist in young mice, n = 4. **B.** Cell parameters recorded in pyramidal neuron in young mice, from left to right: resting membrane potential, rheobase, action potential threshold, and input resistance. n = 4. **C.** Same as in A for adult mice, n = 4. **D.** Same as in B for adult mice, n = 4.

## Notes

### Competing Interest Statement

The authors have declared no competing interest.

